# Investigating side effect modules in the interactome and their use in drug adverse effect discovery

**DOI:** 10.1101/089730

**Authors:** Emre Guney

## Abstract

One of the biggest challenges in drug development is increasing costs of bringing new drugs to the market. Many candidate drugs fail during phase II and III trials due to unexpected side effects and experimental methods remain cost ineffective for large scale discovery of adverse effects. Alternatively, computational methods are used to characterize drug side effects, but they often rely on training predictors based on drug and side effect similarity. Moreover, these methods are typically tailored to the underlying data set and provide little mechanistic insights on the predicted associations. In this study, we investigate the role of network topology in explaining observed side effects of drugs. We find that drug targets are closer in the interactome to the proteins inducing the known side effects of the drug compared to the proteins associated with the rest of the side effects. We show that the interactome based proximity can be used to identify side effects and we highlight a use case in which interactome-based side effect prediction can give insights on drug side effects observed in the clinic.

## Introduction

Drug safety is one of the major driving factors beneath the attrition of drugs, contributing to more than 20% of the clinical trial failures and thus increasing costs associated with drug development [1, 2]. Undesired side effects of drugs are also among the leading causes of mortality in Western countries [3], prompting a clear need for better understanding of drug side effects.

*In silico* association of drugs to side effects offers a cost-effective alternative toward characterizing drug side effects. The first examples of such methods using chemical structure similarity to cluster drugs and predict their potential side effects date back to more than a decade ago [4, 5]. Nonetheless, the large-scale prediction of drug side effects gained immense attention after the availability of SIDER, a resource containing side effect information mined from drug labels [6]. Using the data available in SIDER, several studies sought building prediction models incorporating various similarity measures between drugs and side effects [7, 8, 9]. For instance, Atias and Sharan combined diffusion in side effect similarity network and canonical correlation analysis using chemical fingerprint similarity of drugs to link drugs to side effects [7]. Similarly, Duran-Frigola and Aloy turned to building decision tree based classifiers using various features such as chemical structure, small fragments, drug targets, functional associations, and pathway annotations over-represented among known drug-side effect associations [8]. They used these classifiers to explain the side effect profiles of drugs observed in SIDER and found that the side effects can be predicted reliably for a relatively small portion of the data set. Huang *et al.*, on the other hand, included the interactors of drug targets in the protein-protein interaction (PPI) network in addition to the structural properties of the drug to train a support vector machine and suggested that integrating PPI improved the prediction accuracy substantially [9].

The topology of the human interactome encodes biologically relevant information that can be used to discover novel drug-disease [16, 17, 18], and drug-side effect [19, 20] relationships. Although, some side effects can be explained by the proteins the drug is intended to target, many side effects likely to originate from the interactions of the drug with off-targets or the interactions between these proteins [21]. To understand the role of protein interactions in drug induced arrhythmias, Berger and colleagues identified the neighborhood of disease associated genes for long-QT syndrome in the PPI network and used this neighborhood to predict drugs likely to have risks for QT-interval prolongation [19]. They calculated a random-walk based score from each protein in the PPI network to known disease genes involved in long-QT syndrome, corresponding to the reachability of the proteins from the known disease genes. They then used this score to define a long-QT syndrome specific interactome neighborhood and to rank the drugs based on the targets falling in this neighborhood. Moreover, Brouwers *et al.,* investigated whether the side effect similarity between drugs could be explained by the closeness of the drug targets in a functional PPI network [20]. They observed that only a minor fraction (6%) of drugs whose targets were direct neighbors in the network shared similar side effects, emphasizing the need for taking the global topology of the network into account.

In this study, we aim to investigate whether the global topology of the human interactome can characterize drug side effects. We first define side effect modules as the drug targets elucidating the side effects and check the network-based distances between side effect modules and drug targets. We show that drug targets are closer to the proteins associated with the known side effects of the drug in the network compared to the proteins associated with the rest of the side effects. We then use interactome based closeness to systematically identify side effects of the Federal Drug Administration (FDA) approved drugs in the DrugBank database. Finally, we demonstrate how the interactome based closeness can be used to predict side effects of tamoxifen that are not listed in SIDER.

## Materials and methods

### Data sets

The drugs used in our analysis were retrieved from DrugBank v4.3 database [22]. For all FDA approved drugs, we extracted drug-protein interactions including drug target, enzyme, transporter and carrier interactions (hereafter we simply refer all these proteins as drug targets). Uniprot ids from DrugBank were mapped to ENTREZ gene ids using Uniprot id mapping file (retrieved on October 2015). The SMILES strings of drugs were also downloaded from DrugBank.

We obtained drug side effect information from SIDER v4 [23], a resource containing side effects extracted from drug labels via text mining and mapped the drug ids to DrugBank ids using the PubChem mapping provided in Drug-Bank. We represented the side effects with their preferred terms reported in SIDER. To avoid including drugs whose side effects are not well characterized, we only considered drugs with at least five side effects in SIDER.

For validation purposes, in addition to SIDER, we used OFFSIDES [24], cataloging clinically significant drug side effects from FDA adverse event reporting system. We parsed the OFFSIDES flat file and mapped the drug ids to DrugBank ids using the PubChem mapping provided in DrugBank as we did for SIDER. Only the side effects with observed medical effect were included in the analysis.

We used the human interactome curated in a recent study [25], containing physical interactions between proteins from TRANSFAC[26], IntAct[27], MINT[28], BioGRID[29], HPRD[30], KEGG[31], BIGG[32], CORUM[33], PhosphoSitePlus[34], as well as several large scale studies [35, 36, 37]. The coverage and confidence of this integrated interaction network has been showed to be superior to interaction networks coming from yeast-two-hybrid or functional association data sets [25, 18]. Following the methodology in these studies, the largest connected component of the network, containing 141,150 interactions between 13,329 proteins, was used in the analysis.

### Defining side effect modules

To identify drug targets that contribute to the side effects, we followed the procedure presented in Kuhn *et al.* [38]. For each side effect and drug target we counted the number of drugs with and without the side effect for which the drug target was a known target versus the number of drugs with and without the side effect for which the target was not a known target. We used Fisher’s exact test to calculate the two sided P-value of the observed occurrence of the target with the side effect as follows: The P-values were then corrected for multiple hypothesis testing using Benjamini and Hochberg’s method. We selected the targets that were below 20% false discovery rate to describe the side effect module. In our analysis, we considered the side effects modules that had at least five targets in the interactome. We note that although the proposed approach is applicable to side effects defined by any number of proteins, we use the side effects with at least five proteins to ensure that the side effects in the analysis can be fairly explained by a group of proteins. We provide the side effect module information and the Jupyter Notebook to replicate the analysis in this study at github.com/emreg00/proxide.

### Characterizing closeness between drug targets and side effect modules

Given a network *G*(*V, E*), we defined the following topological measures to quantify the network based closeness between targets of a drug, *T*, and proteins in a side effect module, *S*.

i. *Shortest:* The average pairwise shortest path length between each drug target and side effect module protein.

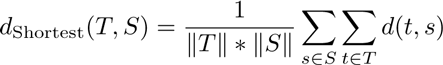

where *d*(*t, s*) is the shortest path length between nodes *t* (a drug target) and *s* (a side effect protein) in the network. To convert the average shortest path length above to a side effect specific z-score for each drug, we normalized *d*_Shortest_(*T, S*) using the mean (*μ*_*d*_Shortest__(*T,S*)) and standard deviation (*σ*_*d*_Shortest__(*T,S*)) of *d*_Shortest_(*T_i_, S*) values calculated for all the drugs *T*_1_*, {T*_2_*, …, T_n_*} in the data set. Accordingly, the closeness between drug *T* and side effect *S* was given by

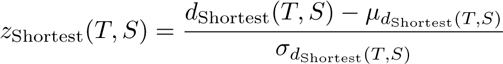

We used Dijkstra’s shortest path algorithm implemented in Python networkx package to calculate the pairwise shortest path length between pairs of proteins in the interactome.
ii. *Closest:* The average shortest path length to the closest protein in the side effect module from the drug targets, given by

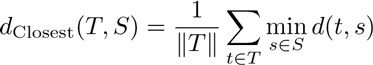

We normalized these values using the mean and standard deviation of the values calculated for all the drugs as it was done above, yielding

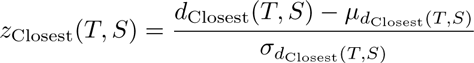
iii. *PageRank:* The average PageRank score of the drug targets when the proteins in the side effect module were used to weight the influence of the nodes in the network. We assigned higher priors to the proteins in the side effect module, 1, compared to the rest of the nodes that were assigned 0.01 and calculated the probability that a random walker in the network would end up in a certain node based on the following formula:

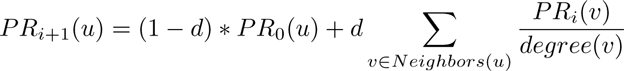

where *u* was the current node in consideration, *v* was a node connected to *u*, *PR_i_*(*u*) was the PageRank score at iteration *i* and *d* is *damping factor* that was set to 0.15. The algorithm was repeated till convergence. The drug - side effect closeness was then defined using

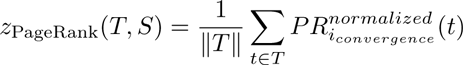

where 
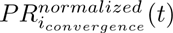
 is the PageRank score of the target *t* normalized using the mean and standard deviation of the PageRank scores of all nodes for the given side effect. We used PageRank with priors implementation in GUILD package [15].
iv. *NetScore:* The average NetScore score of the drug targets when the proteins in the side effect module were used as the source of information passed among the nodes. NetScore scored all the nodes in the network by iteratively propagating the score of the proteins in the side effect module to the neighboring nodes through shortest paths [15]. Unlike conventional shortest path based algorithms, considered the alternative shortest paths in between two nodes, favoring the nodes that were connected with more paths. We used the implementation of NetScore in GUILD software package [15] and initialized the proteins with a score of 1 if they belong to the side effect module and 0.01, otherwise. We limited the number of repetitions the program used to 3 with each iterations in each of them. The drug - side effect closeness was then defined as

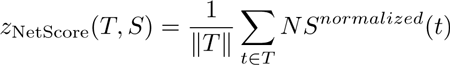

where *NS^normalized^*(*t*) is the NetScore score of the target *t* normalized using the mean and standard deviation of the NetScore scores of all nodes for the given side effect.
v. *Proximity:* The significance of the observed average shortest path length to the closest protein in the side effect module from the drug targets. Interactome based proximity [18] first quantified the average shortest path length between the closest protein in the side effect module and the drug targets (*d*_Closest_(*T, S*) above) and then calculated a z-score corresponding significance of these distances using

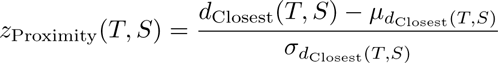

where *μ*_*d*_Closest__(*T,S*) and *σ*_*d*_Closest__(*T,S*) are the mean and the standard deviation of the background distribution of expected minimum shortest path distances between two randomly selected groups of proteins (with the same size and degrees of the original protein sets). The background distance distribution was generated using 1,000 randomly selected protein groups matching drug targets and side effect proteins.

### Drug side effect prediction using network-based closeness

To investigate whether the network-based closeness can predict side effects, for each known and unknown drug and side effect pair, we recorded the five topology based closeness scores (*z_Shortest_*, *z_Closest_*, *z_PageRank_*, *z_NetScore_*, *z_Proximity_*). We then verified whether these topology based scores could discriminate known drug - side effect pairs from the rest by calculating the number of correctly and incorrectly predicted known and unknown drug - side effect pairs at various score cutoffs and checking the area under ROC curve (AUROC) and area under precision-recall curve (AUPRC). The known drug-side effect associations in SIDER and OFFSIDES databases were used as the gold standard positive instances and the remaining associations were assumed to be negative instances. We employed Python scikit-learn package to calculate AUROC and AUPRC values and R for the statistical tests.

## Results

### Side effect modules in the interactome

The available experimental information on the drug targets contributing to the side effects of drugs is often limited to a handful of drug targets [39, 40], hindering a large scale analysis of drug targets inducing the side effects. Alternatively, over-representation analysis of drug targets and side effects can characterize the targets eliciting side effects [38]. Therefore, we define the side effect modules as the groups of drug targets significantly associated with the side effects using the drug target information in DrugBank [22] and SIDER database [23]. Using 1,530 FDA approved drugs and their targets in DrugBank, we identify 1,177 drug target groups associated with the side effects. To confirm that the proteins defining the side effect modules are biologically relevant, we check the overlap between the side effect targets by Lounkine *et al.* [40]. The side effect modules cover at least one protein associated with the side effect for 164 of 241 side effects that are also in the Lounkine *et al.* study. Furthermore, 130 out 265 of the proteins in the identified side effect modules appear among 224 proteins given in the Lounkine data set, covering more than half of the experimentally verified side effect targets.

To understand the interactome based relationship between drug targets and side effect modules, we focus on 537 side effect modules that have at least 5 proteins in the interactome and 817 drugs both known to exert any of these side effects and having at least one target in the interactome. We seek whether topological characteristics of these two groups of nodes, drug targets and side effect module proteins, can explain observed side effects of drugs (Figure 1a). We first turn our attention to the side effect module proteins and ask if the number of proteins in the module or their degree can provide insights on the side effects drugs show. The average module size is 〈*n*_module_〉 = 15.8 among 537 side effects and the largest module, the one of gynaecomastia (enlargement of a man’s breasts), contains 66 proteins. Interestingly, the average degree of all the proteins contributing to a side effect is higher than the average degree of the remaining proteins in the interactome (〈*k*_side effect_〉 = 26.5 vs 〈*k*_non side effect_〉 = 21.1). If the proteins within each side effect module are considered independently, however, the average degree of the proteins in the side effect modules is around the average degree of the interactome (〈*k*_module_〉 = 20.8 vs 〈*k*〉 = 21.2), with peliosis hepatis, an uncommon vascular condition in liver, being the side effect with the highest average degree (〈*k*_peliosis hepatis_〉 = 123.6).

**Figure 1:**
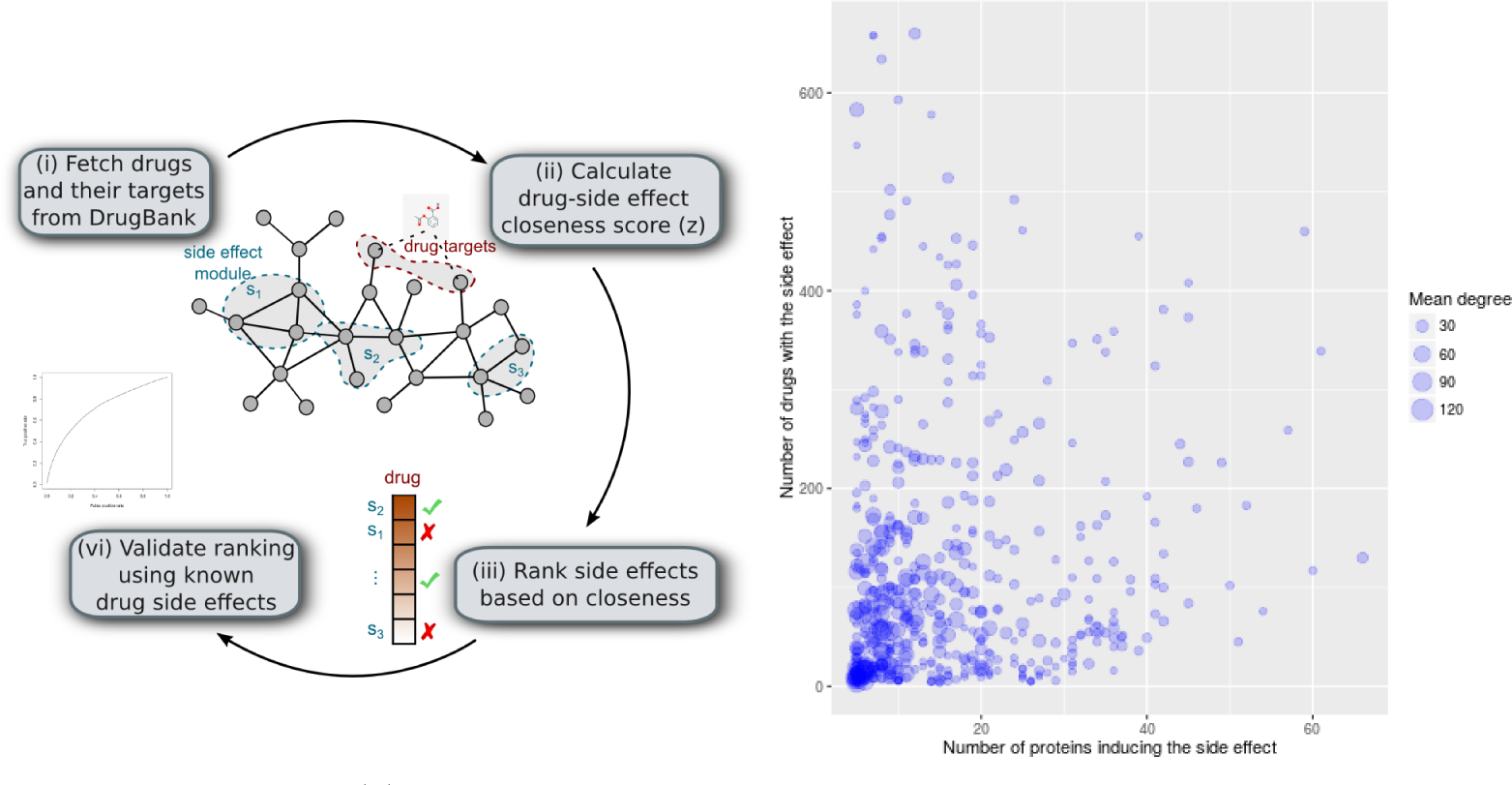
Side effect modules in the interactome and their use in drug adverse effect characterization. **(a)** Schematic overview of the interactome based analysis of drug side effect modules. For each of 817 drugs and 537 side effects, we calculate network based closeness between the drug targets and the proteins inducing the side effect and validate the predictions using known drugside effect associations. **(b)** Each point represents a side effect consisting of proteins identified to be significantly associated to the side effect. The x-axis is the number of proteins in the side effect module and the y-axis is the number of drugs that shows the side effect. The size of the points scales with the median degree of the proteins in the side effect module.

To investigate whether the size and the average degree of the identified side effect modules are higher for the “popular” side effects – the side effects that occur frequently in SIDER–, we look at the number of drugs the side effect is observed and the number and mean degree of the proteins in the side effect module (Figure 1b). The significant but low correlation between the number of drugs showing the side effects and the module size (Spearman’s rank correlation *ρ* = 0.16, *P* = 1.8 × 10^−4^) suggests that the size of the module is not strongly associated to the occurrence of the side effects. On the other hand, the degree of the proteins within the side effect modules is not correlated with the number of drugs the side effect is observed (Spearman’s rank correlation *ρ* = 0.03, *P* = 0.55).

### Network based closeness of drugs and side effects

Next, for each drug and side effect pair in our analysis (817 × 537 pairs), we calculate the network based closeness of the drug’s targets to the side effect module in the interactome using five topological measures (see Methods). We then investigate how well the calculated closeness scores discriminate the observed drug side effects using the known drug side effect associations in SIDER and OFFSIDES databases (Table 1).

**Table 1:**
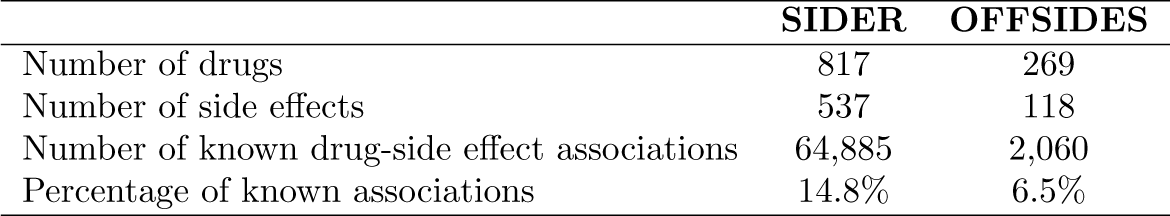
Number of drugs, side effects and known drug - side effect associations included in the analysis according to SIDER and OFFSIDES databases.

|  | SIDER | OFFSIDES |
| --- | --- | --- |
| Number of drugs | 817 | 269 |
| Number of side effects | 537 | 118 |
| Number of known drug-side effect associations | 64,885 | 2,060 |
| Percentage of known associations | 14.8% | 6.5% |

We find that the drugs tend to be closer to the proteins inducing the side effects known to be associated with them compared to the proteins in the rest of the side effect modules (Figure 2). The difference in the closeness values of known and unknown drug - side effect pairs is significant using both SIDER and OFFSIDES side effect associations (one-sided Mann–Whitney U test *P* ≪ 0.05). We observe that NetScore, the method that takes alternative shortest path between drug targets and side effect module proteins and Proximity, the method that compares observed shortest path length between drug targets and the closest side effect module protein to the distances between randomly selected nodes in the network yield a wider range of closeness scores than the remaining methods.

**Figure 2:**
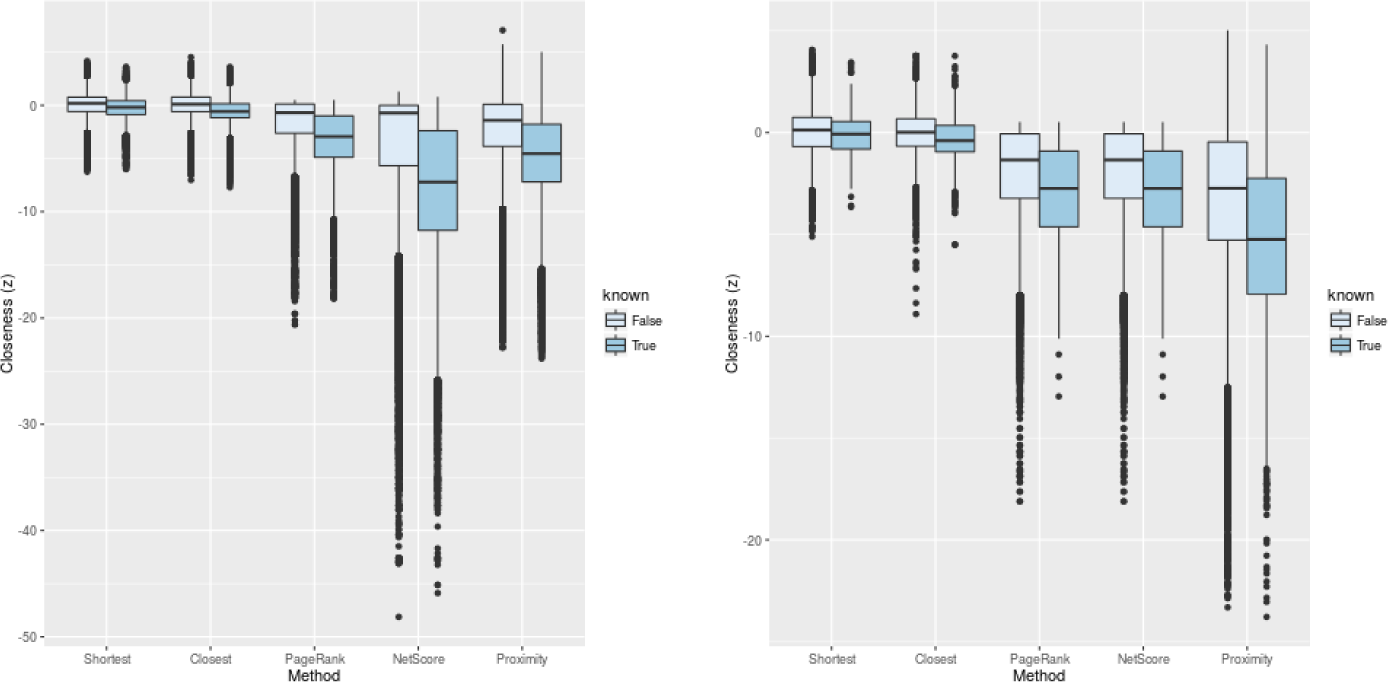
Network based closeness of known and uknown drug - side effect pairs. The closeness between drug targets and side effects calculated using five topological measures (Closest, Shortest, PageRank, NetScore and Proximity) for each of 817 drugs and 537 side effects. Known drug - side effect associations are taken from **(a)** SIDER and **(b)** OFFSIDES.

We then turn to predicting drug side effects using the network neighborhood information of the side effect modules and quantify the closeness between drug targets and side effect modules in the interactome. We use the drug-side effect associations in SIDER and OFFSIDES as the gold standard data to calculate the precision, recall, false positive rate at various closeness score cutoffs and check the area under the ROC curve (AUROC), the area under the precision-recall curve (AUPRC) and the percentage of the drugs for which the highest scoring prediction is a known side effect (Table 2). We see that, overall, the best performing methods are NetScore and Proximity, showing higher prediction accuracy on both SIDER and OFFSIDES data sets compared to the rest of the methods.

**Table 2:** AUROC, AUPRC and percentage of correctly predicted highest ranked drug - side effect pair for various network based closeness methods using SIDER and OFFSIDES associations.

|  | AUROC (%) |  | AUPRC (%) |  | Correct at top (%) |  |
| --- | --- | --- | --- | --- | --- | --- |
|  | SIDER | OFFSIDES | SIDER | OFFSIDES | SIDER | OFFSIDES |
| Shortest | 59.8 | 53.9 | 17.8 | 7.1 | 15.9 | 8.2 |
| Closest | 67.9 | 57.7 | 27.6 | 8.5 | 79.6 | 28.6 |
| PageRank | 69.0 | 59.6 | 27.0 | 8.6 | 55.8 | 13.0 |
| NetScore | 71.7 | 61.9 | 28.8 | 9.6 | 52.1 | 14.5 |
| Proximity | 71.1 | 63.6 | 32.8 | 11.4 | 56.7 | 11.5 |

Despite using only the network topology the AUROCs for NetScore and Proximity scores on SIDER drug-side effect associations are 71.7% and 71.1%, respectively, suggesting that closeness of drugs to side effect modules is predictive of the drug’s adverse effects. We also examine the area under precision-recall curve (AUPRC) and find that NetScore and Proximity achieve AUPRC values of 28.8% and 32.8%, respectively. Furthermore, for 52.1% and 56.7% of the drugs used in the analysis, the highest scoring side effect identified by NetScore and Proximity is reported in SIDER, showing that drug-side effect module closeness can provide insights on the side effects of drugs. On the other hand, when the drug-side effect associations in OFFSIDES database is used, the AUROC drops to 61.9% and 63.6% for NetScore and Proximity, still substantially higher than that would be expected from a classifier producing random predictions (50%). Moreover, only for around 10% of the drugs, the highest scoring side effect is in OFFSIDES, an observation we attribute to the lower coverage of known side effects in OFFSIDES database (6.5%) compared to the SIDER (14.8%, Table 1). Accordingly, due to the higher coverage of drugs and side effects, and better prediction accuracy, in the rest of the text, we use SIDER drug - side effect associations as the gold standard.

### Assessing the effect of the data incompleteness

The current knowledge on drug-target interactions represent only a partial view of the possibly many proteins involved in drug’s action [41]. To account for the potential implications of incompleteness of the drug target data, we analyze the prediction performance of each method on various subsets of drugs and side-effects categorized with respect to the number of drug targets (*m*) and side effect proteins (*n*). Figure 3 shows the AUROC and AUPRC values (*i*) on the original data set containing 817 drugs with at least one target and 537 side effect modules of at least five proteins (*m* ≥ 1, *n* ≥ 5) and when we repeat the analysis using (*ii*) 428 drugs and 537 side effects with at least five targets and proteins (*m* ≥ 5, *n* ≥ 5), (*iii*) 428 drugs with at least five targets and 322 side effect modules with at least ten proteins (*m* ≥ 5, *n* ≥ 10), and finally, (*iv*) 176 drugs and 322 side effects with at least ten proteins (*m* ≥ 10, *n* ≥ 10).

**Figure 3:**
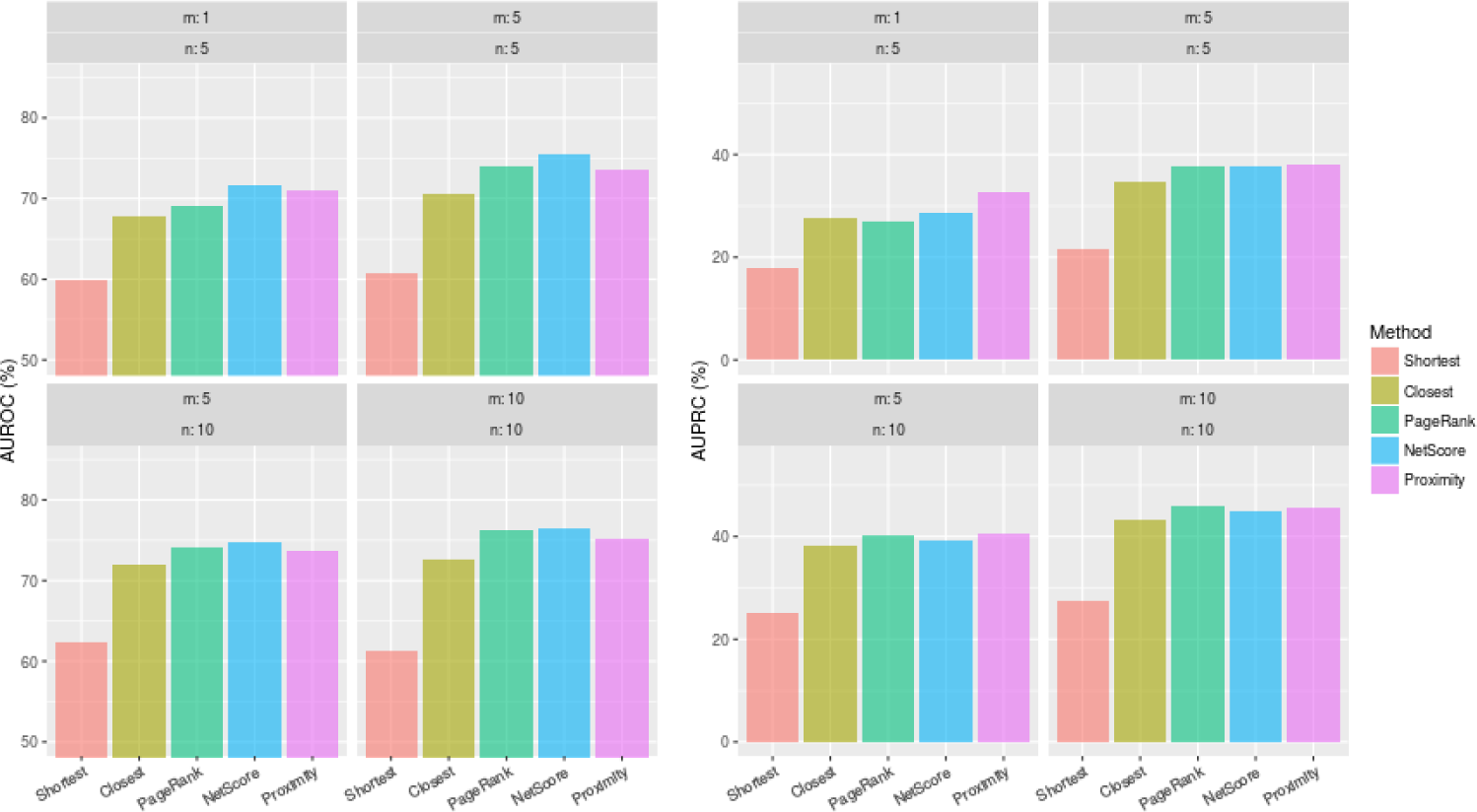
The effect of data incompleteness on prediction performance. The area under **(a)** the ROC curve (AUROC) and **(b)** the precision-recall curve (AUPRC) values when a subset of the drugs and side effects are excluded from the analysis. In each panel, the drugs having less than *m* targets in the network and the side effect modules that have less than *n* proteins in the network are excluded from the analysis.

We find that, as the drugs and side effects associated with more proteins are used, the closeness based predictions improve. Nonetheless, the improvement mainly stems from the higher number of drug targets, as the change in the accuracy is modest when the number of proteins in the side effect modules increases. On the other hand, the AUROC and AUPRC values increase 3-6% when the drugs with more number of targets are used.

### Case study: Top ranking side effects of Tamoxifen

To highlight how interactome based closeness of drug targets can help identifying side effects, we use Proximity, the method that show high overall accuracy according to various performance measures (Table 2). Using only the target information of a given drug, Proximity calculates a network topology based significance of the closeness of the drug to all side effects, allowing us to rank the likelihood of all side effects for any drug with drug target information. Notably, among the drugs in our data set for which the top ranking side effect is not reported in SIDER, we see tamoxifen, an estrogen receptor modulator used for the treatment of breast cancer. Although eight out of ten highest scoring side effects are reported in SIDER, two side effects with very strong association scores, “muscular weakness” and “neuropathy peripheral” are not listed in SIDER (Table 3). We find out that the muscle weakness is indeed a known side effect according to the drug information in Medlineplus (nlm.nih.gov/medlineplus/druginfo/meds/a682414.html). Furthermore, while not indicated in neither SIDER nor Medlineplus, the peripheral neuropathy appears to be a clinically relevant condition reported by several patients in message boards (community.breastcancer.org/forum/78/topics/780591, medhelp.org/posts/Breast-Cancer/tamoxifen-and-neuropathy/show/261680, medhelp.org/posts/Breast-Cancer/Can-longer-term-tamoxifen-cause-peripheral-neuropathy/show/1384498).

**Table 3:** Top 10 side effects predicted for tamoxifen using Proximity.

| Rank | Side effect | in SIDER | Proximity score (z) |
| --- | --- | --- | --- |
| 1 | muscular weakness | 0 | -12.9 |
| 2 | musculoskeletal discomfort | 1 | -12.3 |
| 3 | alopecia | 1 | -12.1 |
| 4 | neuropathy peripheral | 0 | -12.1 |
| 5 | drug interaction | 1 | -11.7 |
| 6 | hepatitis | 1 | -11.7 |
| 7 | diarrhoea | 1 | -11.7 |
| 8 | myalgia | 1 | -11.6 |
| 9 | injury | 1 | -11.5 |
| 10 | discomfort | 1 | -11.3 |

The Proximity score of Tamoxifine to the 14 proteins associated to peripheral neuropathy is *z* = −12.1, suggesting that the drug targets are highly proximal to the side effect proteins in the interactome as a group. This is largely due to seven enzymes (*CYP1A2, CYP2C19, CYP2C8, CYP2C9, CYP2D6, CYP3A4, CYP3A7*) and two transporters (*ABCB1, ABCC2*) tamoxifen is known to bind are in the side effect module. Furthermore, protein encoded by *KIT* gene in the side effect module, is known to be inhibited via phosphorylation by Protein kinase C protein family, a family of proteins targeted by tamoxifen, contributing to the observed proximity to the peripheral neuropathy.

## Discussion

Most existing approaches rely on existing drug side effect associations to predict drug side effects, hindering both the interpretability of predicted associations and the ability to discover novel side effects. In contrast, in this study, we investigate the network based closeness of drug targets to the proteins likely to induce the side effects to explain the observed drug adverse effects. We use the interactome based closeness to predict side effects associated with a drug, providing a mechanistic explanation of the predicted association.

We start with defining the proteins inducing side effects and show that the proteins used to define side effect modules have fair coverage of known side effect inducing proteins. We find that though the proteins likely to induce the side effects show a slight tendency to have higher degrees in the interactome, the effect of degree is not prominent when the side effect modules are considered individually. We also find that the size and the average degree of the identified side effect modules are not higher for the side effects that occur frequently in SIDER. Taken together these findings suggest that the number or degrees of the proteins in the modules are not a good descriptor of observed side effects.

The AUROC values for drug adverse effect prediction reported in the literature range between 60-90% depending on the validation scheme, data sets, predictive models and features (see [42] for a recent review). In particular, compared to the predictor combining canonical correlation analysis using chemical similarity and network diffusion on side effect similarity network by Atias and Sharan, Proximity identifies a known side effect as the top scoring side effect for 56.7% of the drugs in contrast to 34.7% of the drugs reported in the original study [7]. In another study, Huang *et al.* reported an AUROC value (70%) similar to that of using Proximity for the support vector machine based predictor that combined various features including PPI network neighborhood of drug targets and drug structural properties [9].

One drawback of network based methods is that they require that at least a drug target known to interact with a protein in the interactome. Furthermore, they can only be applied to side effects for which a set of proteins inducing the side effect can be identified. Yet, we show that interactome based closeness can systematically detect side effects of 817 FDA approved drugs in DrugBank without relying on the known drug-disease associations. Moreover, network based closeness offers an important advantage over widely used similarity based methods by providing interactome-based insights on the likelihood of a drug to induce a given side effect.

## Acknowledgements

The author is grateful to Dr. Patrick Aloy for providing computational resources for this study and the members of the lab for fruitful discussions. EG is supported by EU-cofunded Beatriu de Pinós incoming fellowship from the Agency for Management of University and Research Grants (AGAUR) of Government of Catalunya.

